# Cerebellar Purkinje cell simple spike activity in awake mice represents phase differences between oscillations in medial prefrontal cortex and hippocampus

**DOI:** 10.1101/173849

**Authors:** Samuel S. McAfee, Yu Liu, Roy V. Sillitoe, Detlef H. Heck

**Author notes:** These authors contributed equally (Samuel S. McAfee, Yu Liu). <u>Corresponding author</u>: Detlef Heck, Ph.D., University of Tennessee Health Science Center, Dept. Anatomy & Neurobiology, 855 Monroe Ave., Memphis, TN 38163.

## Abstract

The cerebellum has long been recognized for its role in tasks involving precise timing, particularly the temporal coordination of movements. Here we asked whether cerebellar might be involved in the temporal coordination of the phases of neuronal oscillations in the medial prefrontal cortex (mPFC) and dorsal hippocampus CA1 region (CA1). These two structures and the cerebellum are jointly involved in spatial working memory. The phases of oscillations in the mPFC and CA1 have been shown to reach a stable alignment (coherence) during the decision making process in a spatial working memory task. Here we report that PC simple spike activity in the cerebellar lobulus simplex in awake, head-fixed mice represents specific phase differences between oscillations in the mPFC and CA1. Most PCs represented phase differences in more than one the conventional frequency bands (delta, theta, beta and gamma). Between the 32 PCs analyzed here, phase differences in all frequency bands were represented. PCs representing phase differences in the theta and low gamma bands showed significant population preference for mPFC phase leading CA1 phase. These findings support the possibility of a cerebellar involvement in the temporal coordination of phase relationships between oscillations in the mPFC and CA1.

## Introduction

The cerebellum was long perceived as a structure exclusively devoted to coordination of movements, but it is now widely recognized for its involvement in cognition, in both experimental animal models and humans 1-4. A neuronal mechanism for cerebellar involvement in cognition has not yet been identified. While the performance of cognitive tasks does not require solving the mechanical problems associated with coordinating inertial masses during body and limb movements, we argue that the precise temporal coordination of neuronal activity is a requirement common to both motor 5-8 and non-motor tasks 9-14. Here we hypothesize that the neuronal mechanisms of cerebellar involvement in cognitive function involves the temporal coordination of neuronal activity by the cerebellum. We asked specifically whether the temporal relationship of oscillatory phases in the neuronal activity in the medial prefrontal cortex (mPFC) and the hippocampal CA1 region (CA1) is reflected in cerebellar Purkinje cell (PC) activity.

The mPFC and CA1 are jointly involved in spatial working memory (SWM) performance. Recent studies have implicated the cerebellum in spatial orientation 15-17, including SWM tasks 17-21. Imaging studies have shown that performance of a SWM task leads to increased activity in several areas of the cerebellum 22. The precise location of cerebellar areas activated during SWM varied between subjects, but the most consistent activation across subjects was seen in the lobulus simplex 22. Taken together, this evidence suggests a joint involvement of the mPFC, CA1 and the cerebellar lobulus simplex in the performance of SWM tasks.

There is broad consensus that the cerebellum plays a key role in several tasks involving precise timing, including the temporal coordination of movements 6,23-29, the perception of temporal patterns 30-34 and predictive timing related to learning 35-37. A form of precise temporal coordination associated with cognitive function is the task-related modulation of coherence of local field potential (LFP) oscillations between two structures 38. An increase in the coherence of LFP oscillations has been suggested as a mechanism to enhance neuronal communication 38,39. The mPFC and dorsal CA1 region express increased coherence of LFP theta and gamma band oscillations during the decision-making process in a SWM task 40,41. Adjusting the coherence of oscillations between two structures requires the precise temporal alignment of the phases of oscillations in the two structures. Here we asked whether the cerebellum is involved in this timing task, targeting specifically the coordination of oscillation phases between the mPFC and CA1.

To address the question whether the cerebellum might be involved in the temporal coordination of phase relationships between oscillations in two structures that are devoted to non-motor brain functions, we performed simultaneous recordings of LFP oscillations in the mPFC, dorsal CA1 and single unit PC spike activity in cerebellar lobulus simplex. Our results show that the phase relationship between oscillations in the mPFC and CA1 are represented in lobulus simplex PC activity. PC phase difference representations were found in delta, theta, beta, low and high gamma frequency oscillations, with different PCs representing different frequency bands. PCs representing phase differences within the theta and low gamma bands stood out, as they all shared a preference for representing phase differences with the mPFC phase leading CA1 phase.

Our findings are consistent with the idea that the cerebellum is involved in the temporal coordination of phase relationships between oscillations in non-motor brain areas and hence that it could have a role in to the coordination of functional connectivity by modulating coherence.

## Results

### Link between PFC-CA1 phase differences and spike rate in individual PCs

We obtained recordings of LFP activities in the mPFC and dorsal CA1 region while simultaneously recording PC simple spike activity in the cerebellar lobulus simplex (Fig. 1). All recordings were performed in awake, head-fixed mice at rest. In order to determine whether the simple spike activity of PCs in lobulus simplex is correlated with phase differences between LFP oscillations in the mPFC and CA1, we quantified PC spike activity as a function of phase difference between oscillations in the mPFC and CA1 for frequencies between 0.5 and 100 Hz. The relationship between spike rate and phase difference was quantified for each frequency band by calculating a vector with direction corresponding to a preferred mPFC-CA1 phase difference and length representing how strongly spike activity reflected that particular phase difference (Fig. 3). Statistical significance of spike modulation as a function of mPFC-CA1 phase difference was determined using bootstrap statistical methods (see Methods section for details). Significant firing rate modulation with mPFC-CA1 phase differences was found in 29 out of 32 PCs (90.6%).

**Figure 1:**
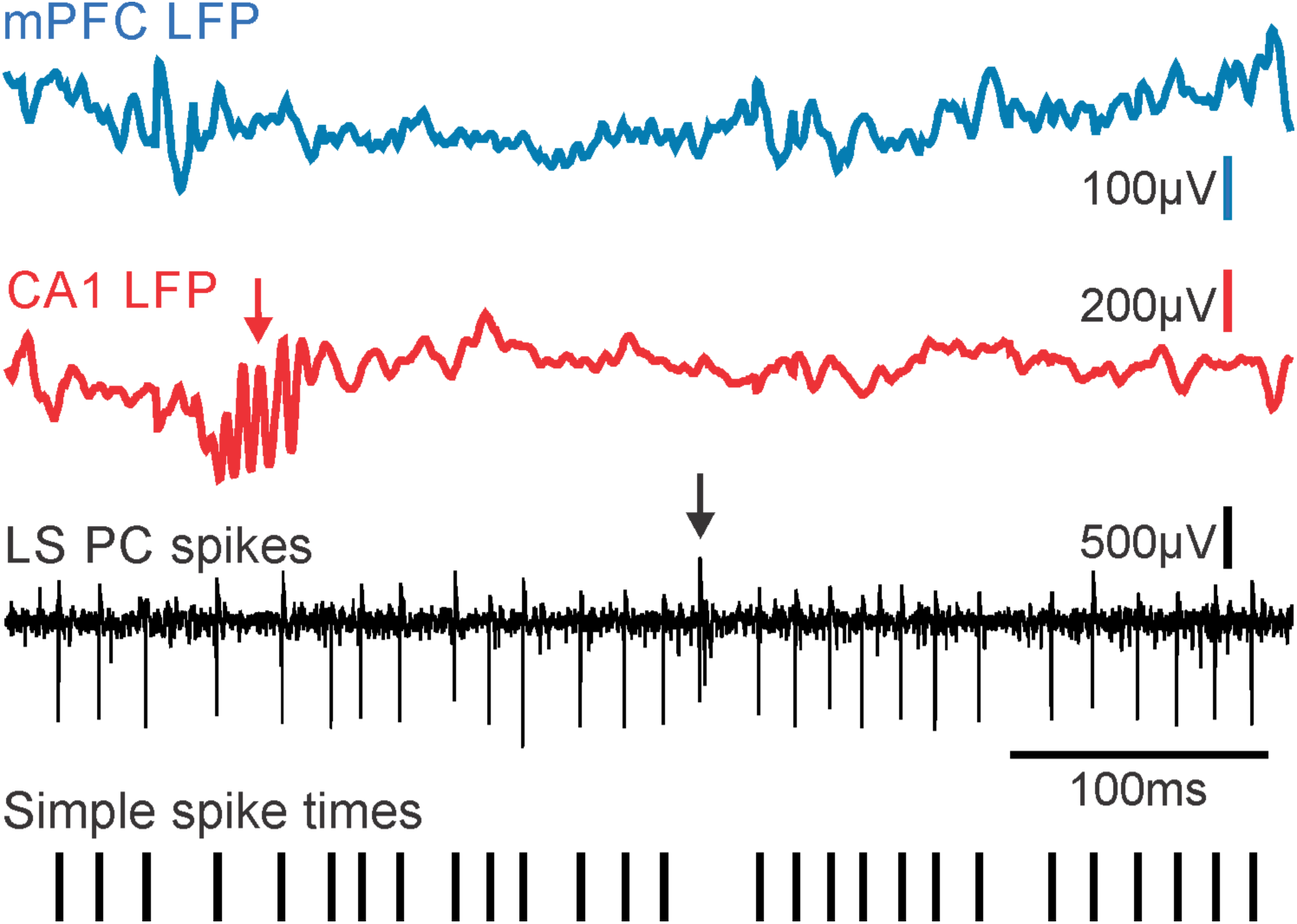
Example of raw LFP signals recorded in the mPFC (blue trace) and CA1 (red trace), and of single unit PC spike activity recorded in the cerebellar lobulus simplex (black trace). The red arrow in the LFP recording from the CA1 region points to a sharp wave ripple event, a brief high-frequency oscillation characteristic for the hippocampus. The presence of sharp wave ripples in the LFP recordings was used for online verification of electrode tip placement within CA1. In the trace of raw PC spike activity a black arrow marks the occurrence of a complex spike, which are characteristic for PCs and were used for online identification of PC activity. The bottom trace shows tick marks representing the time sequence of simple spikes extracted from the raw trace and used for subsequent analysis.

**Figure 2:**
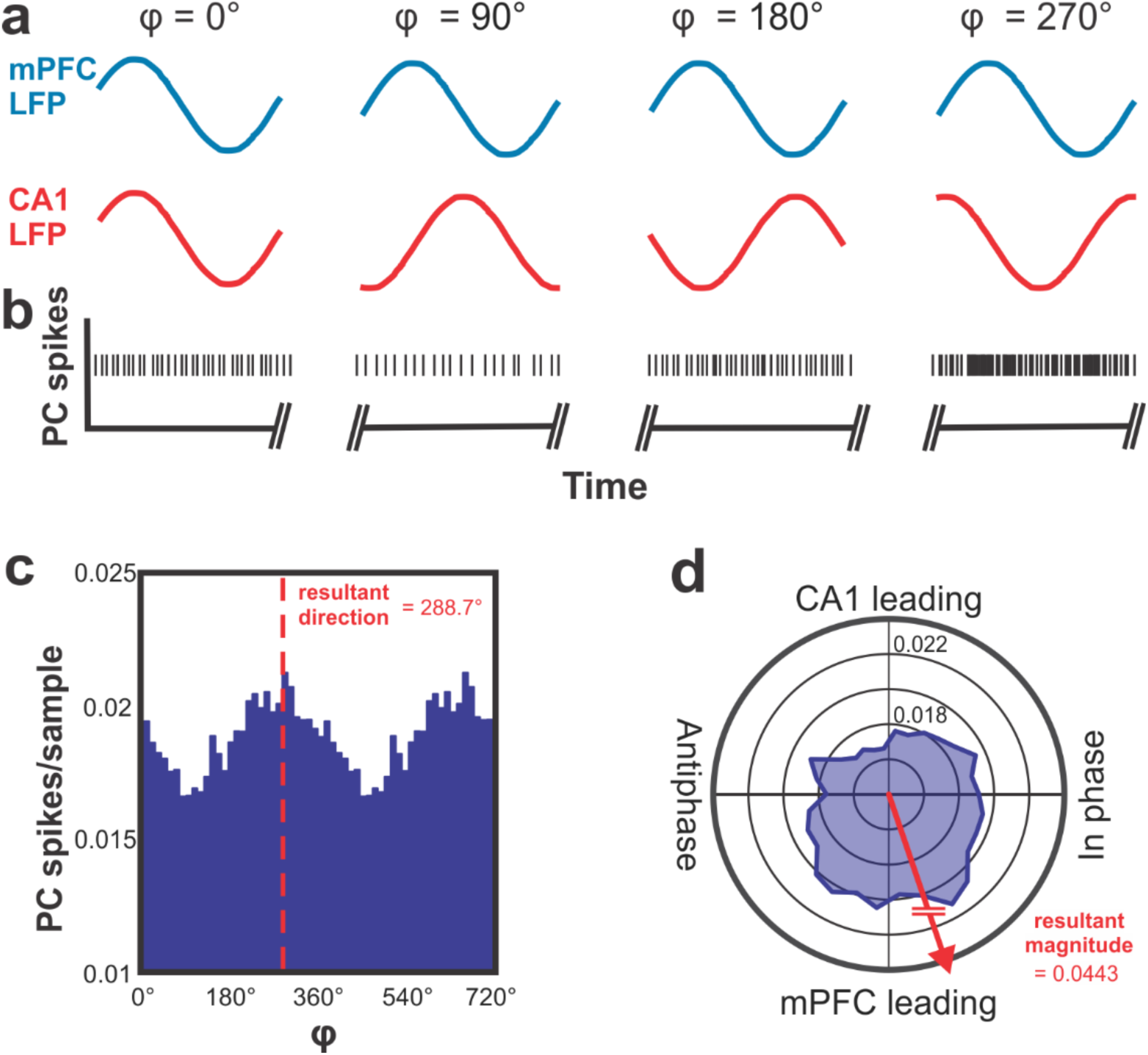
Conceptual illustration of the data analysis applied to determine the correlation between phase differences between mPFC-CA1 oscillatory LFP activity and PC simple spike activity in LS. a) Illustration of hypothetical oscillations at a specific frequency occurring simultaneously in the mPFC (blue traces) and CA1 (red traces) and displaying different phase relationships (φ) at different times. The phase relationship φ is defined as the phase difference relative to the mPFC oscillation. b) Hypothetical PC spikes recorded simultaneously with the LFP activity in the mPFC and CA1 shown in a. The rate modulation of this hypothetical PC shows a significant increase in spike firing when the phase difference between mPFC and CA1 oscillations reaches values around 270°. c) Peri-phase time histogram of real PC simple spike activity. The histogram shows spike activity as a function of mPFC-CA1 phase differences at 11 Hz. The simple spike activity of the PC in this example was significantly modulated as a function of mPFC-CA1 phase difference, with a preference of 288.7°. d) Same data as c represented in polar coordinates. Vectors composed of the angular value ϕ and magnitude of spikes/sample were summated to determine the angular preference of PC activity. The resultant vector magnitude was taken to quantify the degree of modulation, and tested against surrogate results for statistical significance.

**Figure 3:**
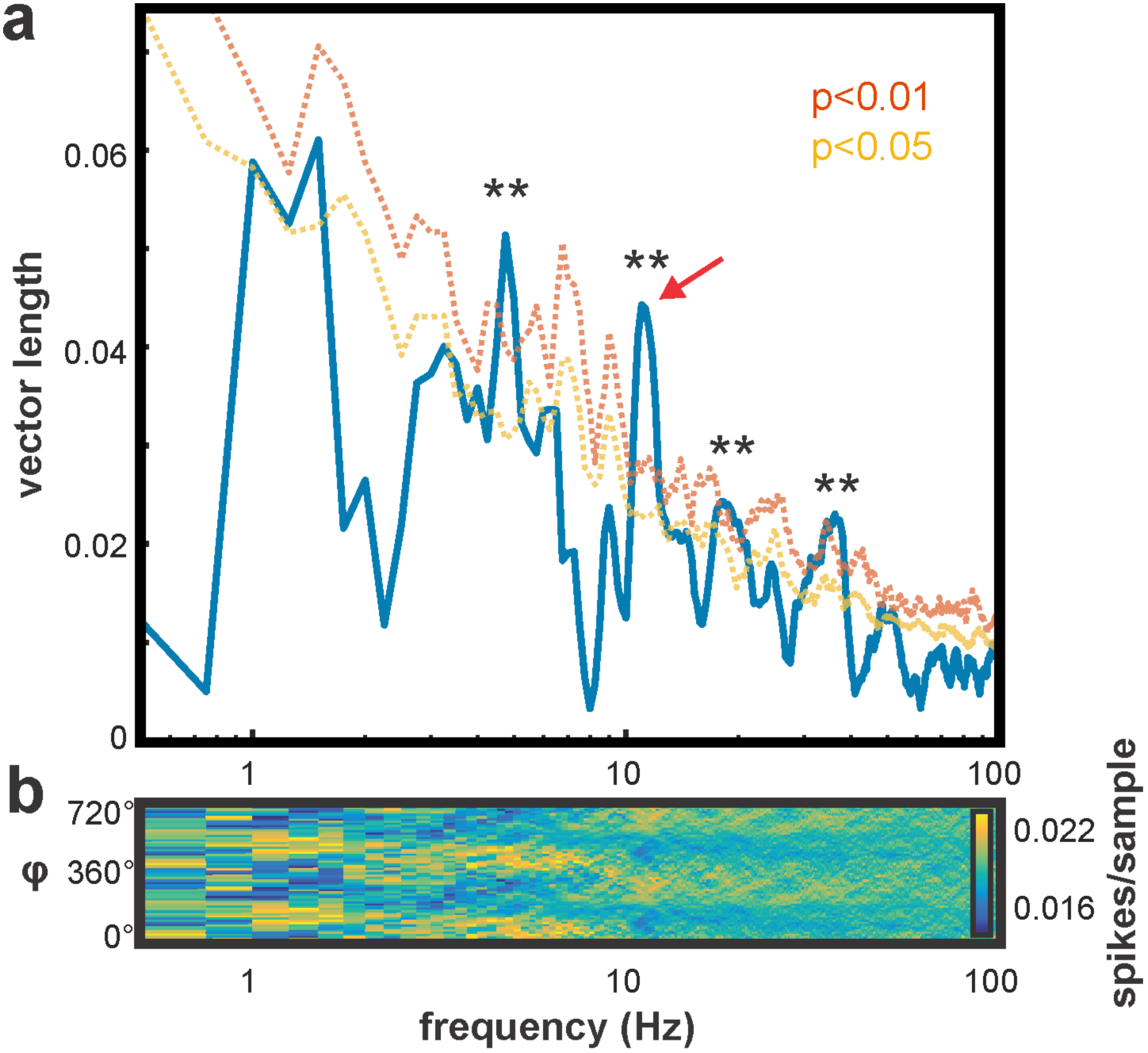
Example analysis for one PC showing how phase differences across the full frequency spectrum of LFP oscillations (0-100 Hz) are represented in this cell’s simple spike activity. a) Blue line shows depth of PC spike modulation, as measured by the resultant vector magnitude (cf. Fig. 2c,d). Significance cutoffs represent the 95%ile and 99%ile boundaries, respectively, of the surrogate result distributions for each frequency obtained by shifting the PC spike recording in time relative to LFPs 200 times. Asterisks indicate frequencies with significant modulation (p<0.01). Red arrow indicates the frequency analyzed to create the histogram and polar plot shown in Fig. 2c,d. b) Pseudocolor plot shows PC spike density as a function of phase difference (ϕ) between mPFC and CA1 LFP oscillations across frequencies between 0.5 and 100 Hz. Analysis was performed within discrete frequency bands of 0.25 Hz width.

### PC activity modulation occurs across the frequency spectrum of PFC/CA1 oscillations

In an analysis of group data we asked what fraction of PCs showed significant phase-related rate modulations at each analyzed frequency (Fig. 4). This analysis revealed that spike rates were most commonly modulated by phase relationships in the theta band, with a maximum at 5 Hz. This did not appear to be a consequence of low frequencies generally being more likely to give a positive result, since fewer significant results were seen at frequencies below 5Hz. Significant results also clustered into modes within each of the other conventional frequency bands: delta (2Hz), beta (17.5Hz), low gamma (35Hz) and high gamma (55Hz).

**Figure 4:**
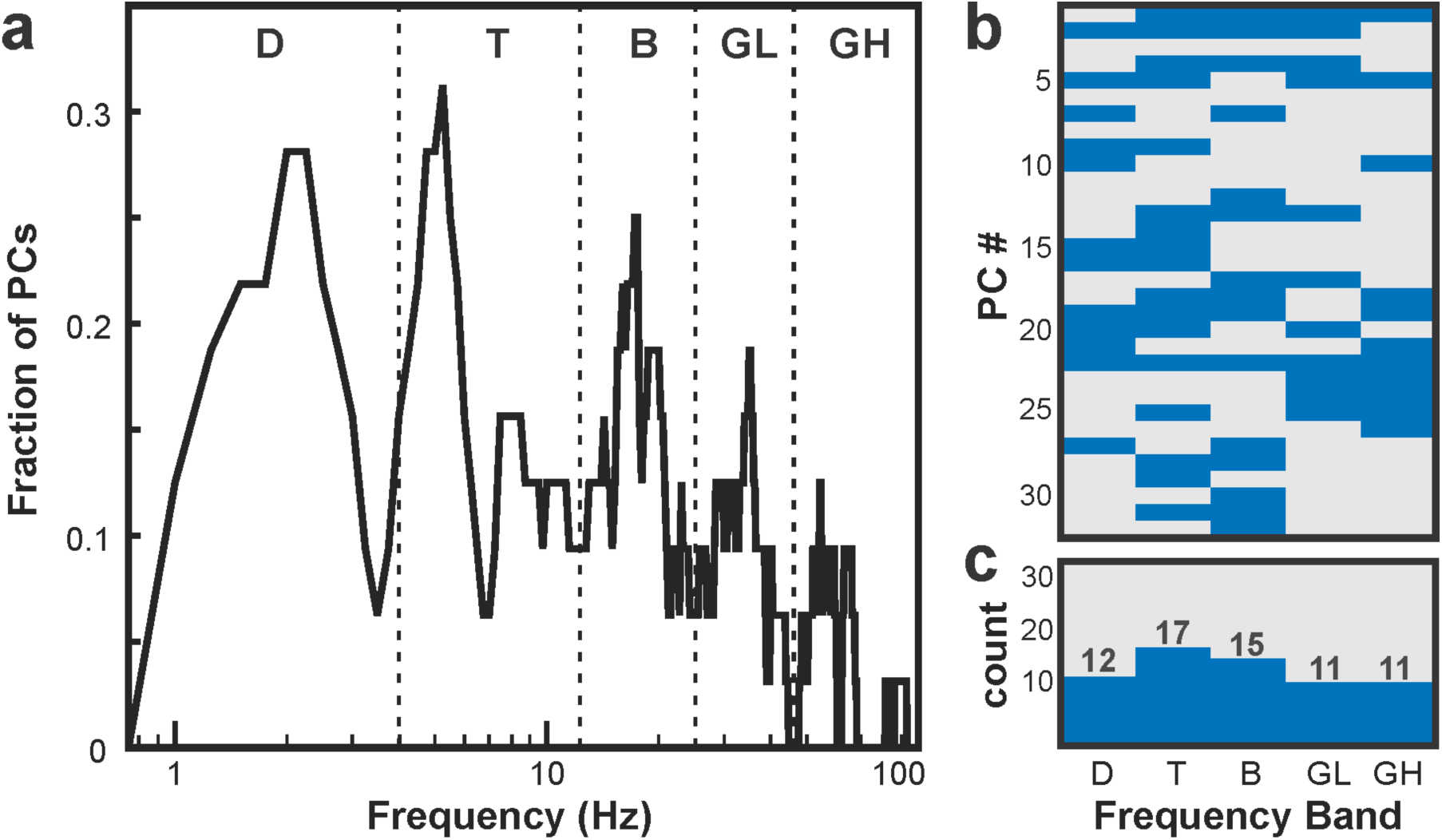
Summary of group results across all analyzed Purkinje Cells (n = 32). a) Plot depicting the fraction of PCs with significantly modulated simple spike activity (p<0.01) within each 0.25 Hz frequency step between 0.5 and 100 Hz. Modal peaks were observed within each of the conventional frequency bands: Delta (D; 0.5-4Hz), Theta (T; 4-12Hz), Beta (B; 12-25Hz), Low Gamma (GL; 25-45Hz), and High Gamma (GH; 45-100Hz). b) Psuedocolor matrix with rows representing individual PCs and columns representing frequency bands. Blue fields indicate a significant correlation between the simple spike activity of the individual PC and phase differences between mPFC-CA1 oscillations in the specific frequency band (p<0.01). Grey fields indicate no significant correlation. c) Histogram showing the number of PCs with significant spike-phase difference correlations within each frequency band.

### PC modulation in low gamma and theta may be linked

Many PCs (68.7%) displayed a significant phase-difference dependent modulation of spike activity in multiple frequency bands (e.g. Fig. 3 for individual example; Fig. 4 b for group data). To investigate whether modulation across the different frequency bands could be linked, we calculated the conditional probability for each frequency band combination (Fig. 5). This analysis revealed a peak in conditional probability between low gamma and theta oscillations, suggesting that a sub-population of PCs might represent phase-differences between oscillations in these two frequency bands either simultaneously or at different times.

**Figure 5:**
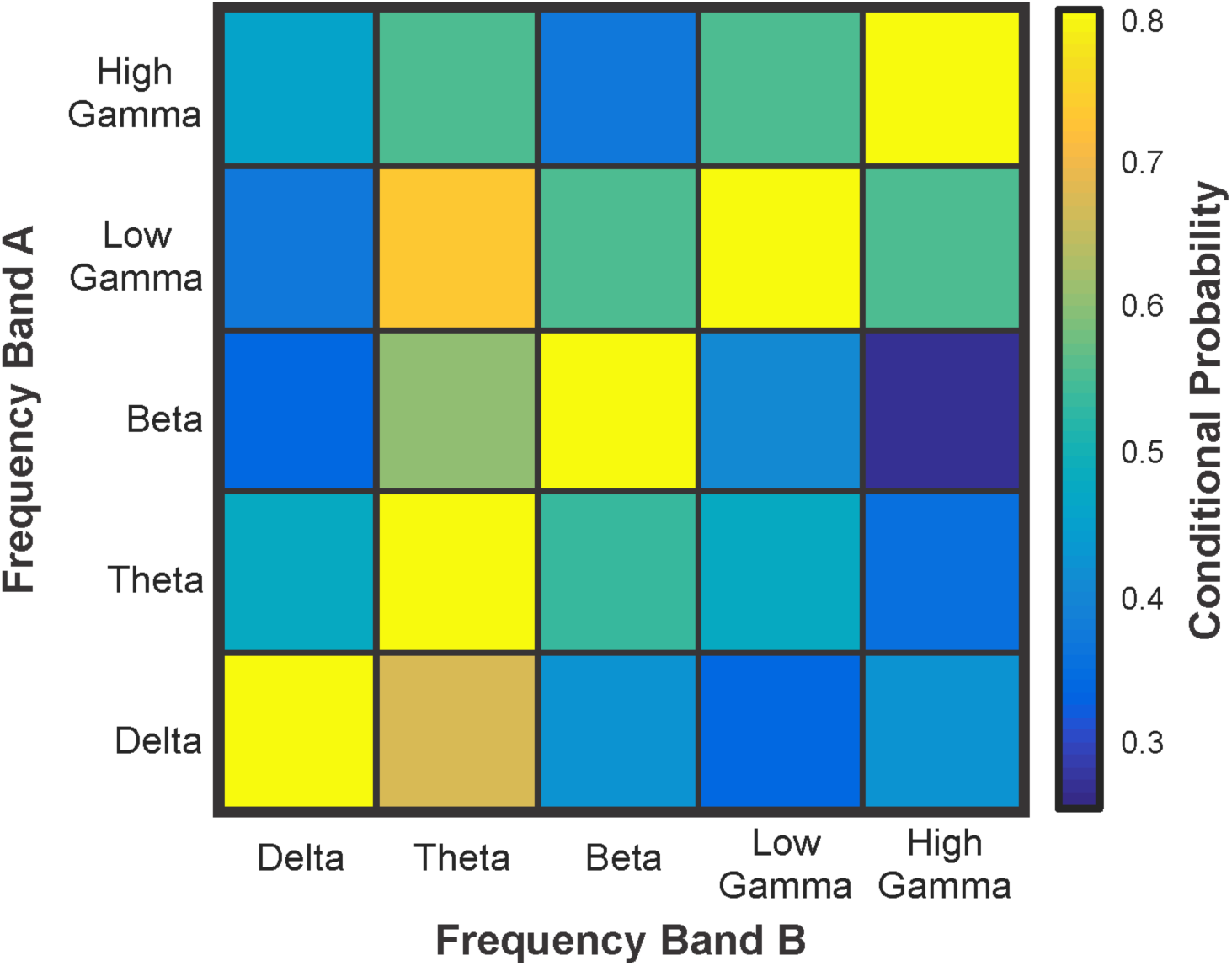
Conditional probability matrix showing the probability of PCs with simple spike activity correlated to phase differences in one frequency band (Band A) to also have a correlation with phase differences in a second frequency band (Band B). Colors represent conditional probability. The highest conditional probability existed for PCs representing phase differences in the Low Gamma (25-45Hz) band, which had a >70% probability to also represent phase differences in the Theta (4-12Hz) band.

### Group-wise phase relationship preferences of PC activity

We further investigated whether significantly modulated PCs showed consistent mPFC-CA1 phase relationship preferences in each frequency band. To do this, we compared the directional values of the vectors at the frequency of greatest spike modulation (Fig. 6). PCs representing phase differences in the theta and low gamma bands showed a significant preference, i.e. generated the highest simple spike activity, for phase differences corresponding to mPFC phase leading CA1 phase (Fig. 6 b,d). PCs representing beta band phase differences showed a clear trend towards a preferred mPFC leading phase difference (Fig. 6 c).

**Figure 6:**
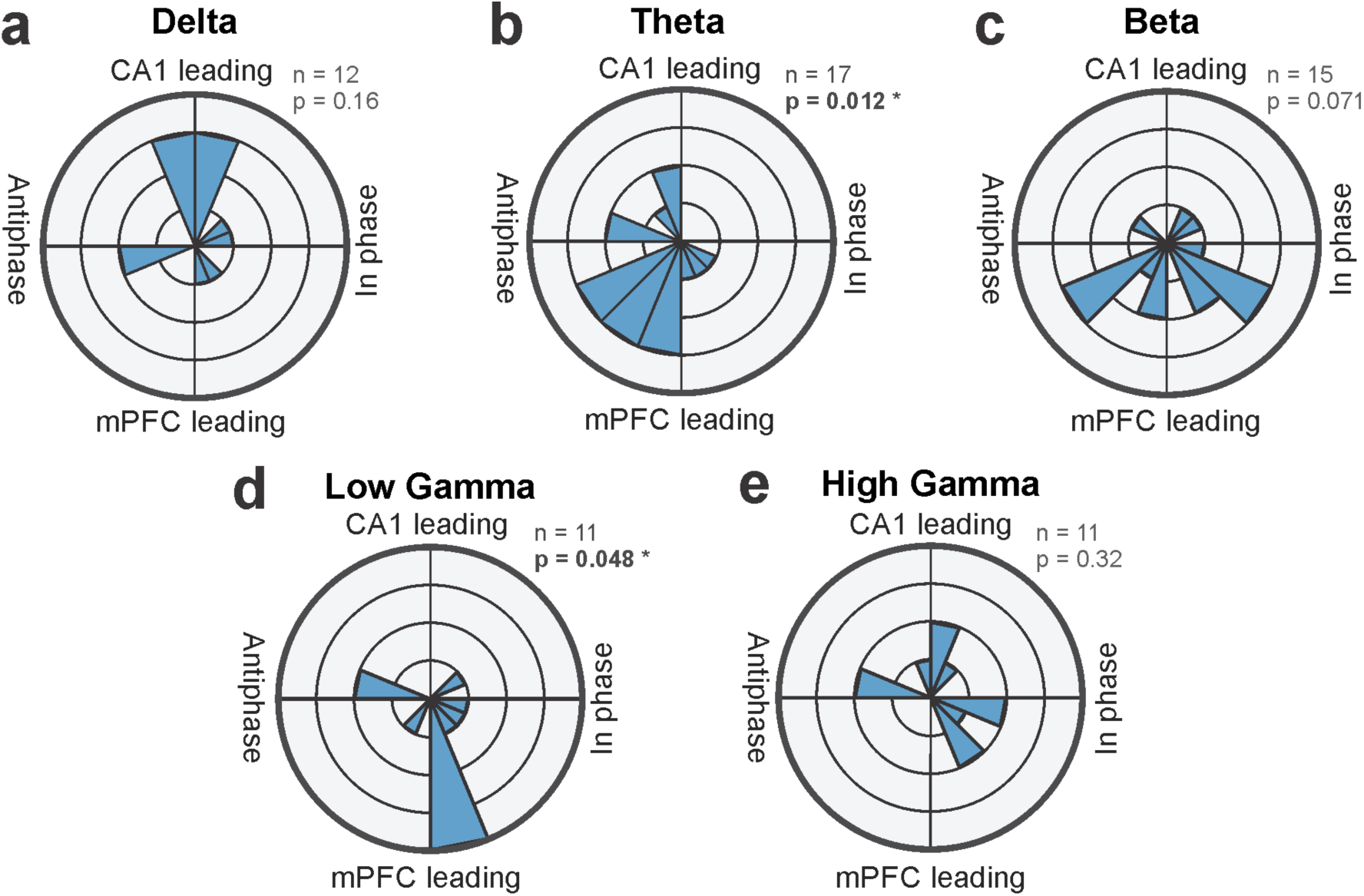
Analysis of phase differences values of mPFC-CA1 oscillations associated with maximal simple spike activity within each conventional frequency band. Only PCs with significant spike-phase-difference correlations within each frequency band were included in the analysis. The number of PCs is indicated next for each polar plot. Probabilities were calculated using a Rayleigh test. Resulting p values are shown next to each polar plot: a) delta, b) theta, c) beta, d) low gamma, e) high gamma. PCs with significant spike-phase-difference correlations in the theta (b) and low gamma (d) frequency bands showed a significant population preference for phase differences with mPFC oscillatory phase leading CA1 phase.

## Discussion

Here we report that simple spike activity of lobulus simplex PCs in awake, head fixed mice represents phase differences between LFP oscillations in the mPFC and CA1. We analyzed single unit spike activity of 32 PCs that were recorded simultaneously with LFP activity in the mPFC and CA1. Each PC represented phase differences between oscillations in one or more of the conventional frequency bands between 0.5 to 100 Hz (delta, theta, beta, low gamma and high gamma). Across the group of 32 PCs analyzed here, phase differences in all frequency bands were represented by multiple PCs, with the largest number of PCs representing phase differences in the theta band.

An analysis of conditional probability revealed a potential link between theta and low gamma oscillations in that PCs representing phase differences in the low gamma band had a particularly high probability to also represent phase differences in the theta band. Interestingly, theta and low gamma oscillations stood out again in an analysis of which specific phase difference values were represented in each PCs activity: there was considerable variability among PCs regarding the “preferred” phase difference values they represented. However, PCs representing phase differences in the theta and low gamma bands showed a significant population bias for phase differences corresponding to mPFC phase leading CA1 phase. There was no significant population bias for a preferred phase differences in the other three frequency bands (delta, beta and high gamma), although there was a trend (p = 0.071) for PCs in the beta band towards a population bias for increased activity when mPFC phase was leading CA1 phase.

The results we present here are based on data collected in awake but head fixed mice. While we thus cannot relate our findings to a controlled behavior, the three structures we simultaneously recorded from are known to be jointly involved in spatial orientation and SWM tasks 42-44. The roles of the mPFC and CA1 in memory tasks in general and in SWM in particular, have been widely investigated and documented 45,46.

Evidence for the involvement of the cerebellum in SWM comes from lesion and stimulation studies in rodents 18,47-49 and imaging studies in humans 22,50. While multiple cerebellar areas seem to be involved in SWM, imaging studies have suggested that the lobulus simplex may play a key role 22.

There is increasing recognition of the crucial role of oscillatory neuronal activity, particularly the synchronization or coherence of oscillations in memory function. Relevant to the findings we presented here are numerous studies that have shown that the mPFC and dorsal CA1 region express increased coherence of LFP oscillations in the theta and gamma band during the decisionmaking process in a SWM task 41,51-54. These decision-related coherence increases are considered critical to SWM decision-making 53.

Coherence is a quantitative measure of the stability of the phase relationship or the phase-locking of two oscillations with similar frequencies. Coherence can take on values between 0 and 1. Two oscillations with a perfectly stable phase relationship over time would have a coherence of 1, irrespective of whether there is a phase delay or advance between the oscillations, as long as the delay is stable 55.

Increasing the coherence of LFP oscillations between two brain structures has been suggested as a mechanism to temporarily enhance neuronal communication between specific structures at a particular time in order to facilitate task performance 38,39. This notion seems consistent with findings showing that SWM related increases in mPFC-CA1 coherence reach higher values during correct decisions than during incorrect decisions 41,52,53. The task-dependent modulation of coherence between two brain structures implies the existence of a mechanism that coordinates the phase differences of oscillations in communicating structures.

The neuronal mechanisms that coordinate phase relationships between brain structures are currently unknown. Our findings provide a first indication of a possible role of the cerebellum in coordinating the phase relationships of neuronal oscillations in the mPFC and CA1, and thus a possible cerebellar involvement in the control of mPFC-CA1 functional connectivity via coherence.

How cerebellar PCs come to represent phase differences between mPFC and CA1 oscillations remains to be determined. Is phase difference information already present in mossy fiber input or does the cerebellum receive phase information from both structures with the cerebellar network extracting phase difference information from those inputs? To the best of our knowledge, no structure has been identified that represents mPFC-CA1 phase differences and projects to the cerebellum. Thus, based on current knowledge and because the cerebellum has reciprocal connections with both the mPFC 3,56,57 and the hippocampus 58-60, we consider the latter scenario more likely. This then calls for an explanation for how the cerebellar network extracts phase difference information from mossy fiber input, assuming that the mossy fiber activity carries phase-information about oscillations in each structure. Considering that any phase difference for a given frequency can be represented as a time interval, a possible neuronal mechanism can be derived from the unique cerebellar network architecture. Parallel fibers have been suggested to serve as delay-lines 61 that could be used to transform phase-difference encoding delays between mossy fiber spike events into synchronous inputs to PCs by delaying the synaptic input signaling the phase leading signal and allowing it to exactly coincide with the input signaling the phase lagging signal 62-64.

A third possibility is that cerebellar output causes mPFC and CA1 oscillations to assume a specific phase difference. This could be accomplished if the cerebellum, via the thalamus, provided rhythmic, appropriately phase-delayed inputs to both structures, which, after a few cycles would entrain the oscillations in each structure. Such entrainment of oscillations in one brain structure to rhythmic input from another has been observed in the visual system in the context of attention selection 65, but can also be induced by external influences such as rhythmic trans-cranial current stimulation 66. Finally, all of the above mechanisms may play a role, each possibly linked to different structures, frequencies or tasks.

In summary, our findings reveal a new aspect of cerebellar neuronal activity, which links the cerebellum to phase-relationships between oscillations in two non-motor structures. Provided the increasingly recognized potential of coherence of oscillations as a means for controlling functional connectivity between brain structures 39,67-70, our findings suggest the possibility of a cerebellar involvement in the coordination of neuronal communication through the modulation of coherence.

## Methods

### Animals

Experiments were performed on six adult male C57BL/6J (B6) mice (>8 weeks old, 18–25 g body weight). Male mice were chosen to avoid outcome variability due to menstrual cycle (e.g. 71. Mice were housed in a breeding colony with 12-hour light/dark cycles in standard cages with ad libitum access to food and water. All experiments were performed during the light cycle, between 12:00 noon and 17:00 hours. None of the mice had undergone any previous experimental procedure. All animal experimental procedures adhered to guidelines approved by the University of Tennessee Health Science Center Animal Care and Use Committee. Principles of laboratory animal care (NIH publication No. 86–23, rev. 1996) were followed.

### Preparation for head-fixed recording

In preparation for surgery, mice were initially anesthetized in an induction chamber and anesthesia was induced with 3% Isoflurane in oxygen. Anesthetized mice were transferred to a stereotaxic head frame and anesthesia was maintained with 1-2.5% Isoflurane in oxygen delivered via a nose cone. Isoflurane concentration was controlled with a vaporizer (Highland Medical Equipment, CA), and maintained at the lowest concentration at which mice failed to show reflexive withdrawal to a toe pinch. Core body temperature was measured and maintained at 37°C using a feedback-controlled heating pad with a rectal thermometer (FHC Inc., Bowdoinham, ME).

Details of the surgical procedure to prepare for head-fixed recording have been described elsewhere 72. In brief, scalp fur was removed and the exposed skin treated twice with iodine. The skull was exposed and cleaned and dried with a scalpel and sterile cotton swabs. Two craniotomies were made over the left prefrontal cortex and hippocampus and a third one over the right cerebellar lobulus simplex. The dura was left intact and covered with triple antibiotic ointment (Walgreens Co., US) to maintain moisture and prevent infection. A cylindrical plastic chamber was placed on each craniotomy for mechanical protection. Three small screws were implanted into the skull to serve as anchors for acrylic cement. A head fixation bar was placed onto the skull and held in place with a drop of cyanoacrylate glue. Finally, acrylic cement (Co-Oral-Ite Manufacturing, New Franken, WI) was applied to the skull screws, the plastic chambers and head fixation bar and allowed to cure. Mice were injected with Carprofen (5mg/kg) for analgesia once at the end of surgery before discontinuing anesthesia and again the morning of the next day. Mice were allowed to recover from surgery for 3 to 4 days prior to recording.

### Electrophysiological recordings

During recordings mice were standing on a flat surface, their bodies gently covered with a loosely fitting ½ tube and the head fixation bar was secured to two stationary metal arms using small machine screws. Mice were allowed to adapt to head fixation for at least fifteen minutes prior to recording. Once the mice were adapted, the antibiotic ointment was removed from the chambers and the chambers were rinsed and filled with sterile saline.

A single 7-electrode micromanipulator and headstage (System Eckhorn; Thomas Recording, Germany) was used to record simultaneously from the three brain areas. This system allows micrometer precise control of the penetration depth of each electrode individually. A custom made set of guiding tubes was prepared to guide electrodes into each area. Guiding tubes were lowered into the plastic chambers so that their tips were just above the dura. The extracellular recording electrodes (glass-insulated tungsten/platinum; 80 μm O.D.; impedance, 3–7 M) were then slowly advanced through the dura to their target location. Hippocampal recording electrodes were positioned such that sharp wave ripples (SWRs), a brief high-frequency local field potential (LFP) oscillation characteristic for the hippocampus 73, could be clearly identified (Fig. 1). Cerebellar recording electrodes were advanced into the lobulus simplex, until a single unit spike train with characteristic complex spikes, which identify Purkinje cells was isolated. Recording locations in the prefrontal cortex were selected following stereotaxic coordinates relative to Bregma 74 and were verified anatomically using small electrolytic lesions created at the end of the final recording session (left mPFC: lat. 0.5 mm, ant. 2.8 mm; left CA1: lat. 2.0 mm, ant. -2.3 mm; Cbl. lobulus simplex: lat. 2.0 mm, ant. -5.52 mm).

Recordings were repeated three times on three successive days for each mouse. After completion of each recording session, the chambers were rinsed and re-filled with triple antibiotic ointment and mice were returned to their home cages. At the end of the last recording sessions electrolytic lesions were created by passing a small current (5 μA/10 s) through the recording electrodes.

Within 12 hours of creating the lesion, mice were euthanized and then transcardially perfused. After perfusion, the brain was removed and post-fixed in 4% formaldehyde for 24 h. Brain tissue was cut into 50 μm thick coronal sections, which were mounted and stained with cresyl violet. The location of lesion sites was determined using a stereotaxic atlas of the mouse brain 74.

LFPs and spike signals were separated by band-pass filtering at 0.1 to 200 Hz and at 200 Hz to 8 kHz, respectively, using a hardware filter amplifier (FA32; Multi Channel Systems, Germany). Filtered and amplified voltage signals were digitized and stored on a computer hard disk (16 bit A/D converter; sampling rate, >20 kHz for action potentials, >1 kHz for LFPs) using a CED power1401 and Spike2 software (both Cambridge Electronic Design). This study focused on the relationship between single-unit spike activity in cerebellar PCs and LFP oscillations in the PFC and CA1. After stable LFP recordings were established in PFC and CA1 no further attempt was made to isolate single unit spike activity in the two areas.

### Analysis of PC single-unit spike data

Single-unit spike activity was recorded from 32 PCs located within cerebellar lobulus simplex in 6 mice. PCs were identified based on their location and firing characteristics such as the presence of complex spikes as well as sustained high-frequency simple-spike firing rates 75. Simple spike activity was identified off-line using a shape-based spike-sorting algorithm in the Spike2 software. In our sample of 32 PCs simple spike firing rates ranged from 36.5 to 157.9Hz, with a mean of 89.1 Hz. Complex spikes were excluded from the analysis. Because of their expected low frequency (~1 Hz), the number of complex spikes in our data samples was too low for meaningful statistical evaluation of their relationship with mPFC-CA1 with phase differences.

### Pre-preprocessing of LFPs

Only sections of data with stable single-unit spike isolation and LFP recordings, which were free of movement artifacts, were included in analysis. CA1 recording sites were included only if they showed clear sharp wave ripples (Fig. 1, CA1 LFP). 60Hz noise was diminished using a hum-removal algorithm in Spike2 that subtracted 60Hz components rather than applying a notch-filter, which left the power spectrum of LFP signals intact.

### Estimation of LFP phase relationships and corresponding spikes

In order to determine phase relationships of LFP oscillations in PFC and CA1 over time, LFPs were band-pass filtered for frequencies 0.5-100Hz. FIR filters were applied in frequency steps of 0.25Hz with bandwidth ranging from 0.5-10Hz with increasing frequency. Filter order was determined to be the number of samples within 5 cycles of the center frequency. The Hilbert transform was performed on each band-pass-filtered signal, and instantaneous phase values were defined as the angular value derived from the complex-valued analytic signal. The phase relationship between PFC and CA1 oscillations within each frequency band was determined in terms of the difference in instantaneous phase values, with mPFC phase being subtracted from CA1 phase. The phase relationships were then designated into 30 angular bins of 12° width, and the corresponding spikes within each bin were counted. The spike count in each bin was divided by the total number of angular samples in the bin to return a measure of spike density (Fig. 2 c). Each bin was then treated as a vector with an angular value of the phase relationship and magnitude of the spike density in order to calculate a resultant vector with magnitude reflecting degree of modulation and direction of preferred phase (Fig. 2 d).

### Statistical analysis

Bootstrap statistical analysis of the relationship between spike activity and PFC-CA1 phase differences was performed by calculating vector lengths for 200 surrogate data sets. Each surrogate data set was created by random, circular shifting of the spike sequence. The resulting 200 surrogate vector values were rank-ordered for each frequency and the 95th and 99th percentile values were determined. Real vector values greater than the 95th percentile of the surrogate vector distribution were considered statistically significant, and those greater than the 99th percentile were selected for further group analysis.

In order to determine whether significantly modulated PCs preferred specific mPFC-CA1 phase differences, we tested for consistency across resultant vector directions. For statistical evaluation of phase-difference-specific PC modulation in each frequency band, we applied a Rayleigh test on the directions of vectors at the frequency of greatest spike modulation.

## Acknowledgements

We thank Shuhua Qi for technical assistance and Michael Nguyen for custom machined parts. This work was supported by internal funds from the College of Medicine, the Neuroscience Institute and the Department of Anatomy & Neurobiology of the University of Tennessee Health Science Center.

## Author contributions

D.H.H. conceived the original idea. D.H.H. and Y.L. designed the experiments. Y.L. performed the experiments. S.S.M. developed and performed the data analysis. D.H.H. and S.S.M. wrote the paper. Y.L. contributed to the writing and to data analysis. R.V.S. contributed to the writing.

## Competing financial interests

The authors declare no competing financial interests.

